# Complex dynamics in an SIS epidemic model induced by nonlinear incidence

**DOI:** 10.1101/331678

**Authors:** Ruixia Yuan, Zhidong Teng, Jinhui Li

**Author notes:** These authors contributed equally to this work.

## Abstract

We study an epidemic model with nonlinear incidence rate, describing the saturated mass action as well as the psychological effect of certain serious diseases on the community. Firstly, the existence and local stability of disease-free and endemic equilibria are investigated. Then, we prove the occurrence of backward bifurcation, saddle-node bifurcation, Hopf bifurcation and cusp type Bogdanov-Takens bifurcation of codimension 3. Finally, numerical simulations, including one limit cycle, two limit cycles, unstable homoclinic loop and many other phase portraits are presented. These results show that the psychological effect of diseases and the behavior change of the susceptible individuals may affect the final spread level of an epidemic.

## Introduction

In generic, the population, for an SIS model, is always separated into two compartments, susceptible and infective individuals. And, in most SIS epidemic models (see Anderson and May [2]), the incidence takes the mass-action form with bilinear interactions. However, in practical, to describe the transmission process more realistic, it is necessary to introduce the nonlinear contact rates [1].

Actually, various forms of nonlinear incidence rates have been proposed recently. For example, in order to incorporate the effect of behavioral changes, Liu, Levin, and Iwasa [3] used a nonlinear incidence rate of the form

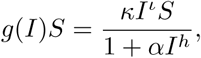

where *κI^ι^* represents the force of infection of the disease, 1/(1 + *αI*^*h*^) is a description of the suppression effect from the behavioral change of susceptible individuals when the infective population increases. *ι, h* and *κ* are all positive constants, and *α* is a nonnegative constant. See also Hethcote and van den Driessche [4], Moghadas [5] and Alexander and Moghadas [6, 7], etc.

In [10], Ruan discuss a specific infection force

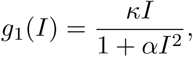

see Fig. 1(a), which can describe the effects of psychology function caused by protection measures and intervention policies. Obviously, *g*_1_(*I*) is increasing with small *I* and decreasing with large *I*, that is, *g*_1_(*I*) is non-monotone. The function *g*_1_(*I*) can be used to interpret the psychological effect: for a very large number of infective individuals, the infection force may decrease as the number of infective individuals increases, since large number of infective may lead to the reducing of the number of contacts per unit time. For example, in 2003, epidemic outbreak of severe acute respiratory syndrome (SARS) had such psychological effects on the general public (see [11]), and aggressive measures and policies had been taken, such as border screening, mask wearing, quarantine, isolation, etc. And also [11] showed that either the number of infective individuals tends to zero as time evolves or the disease persists.

**Fig 1:**
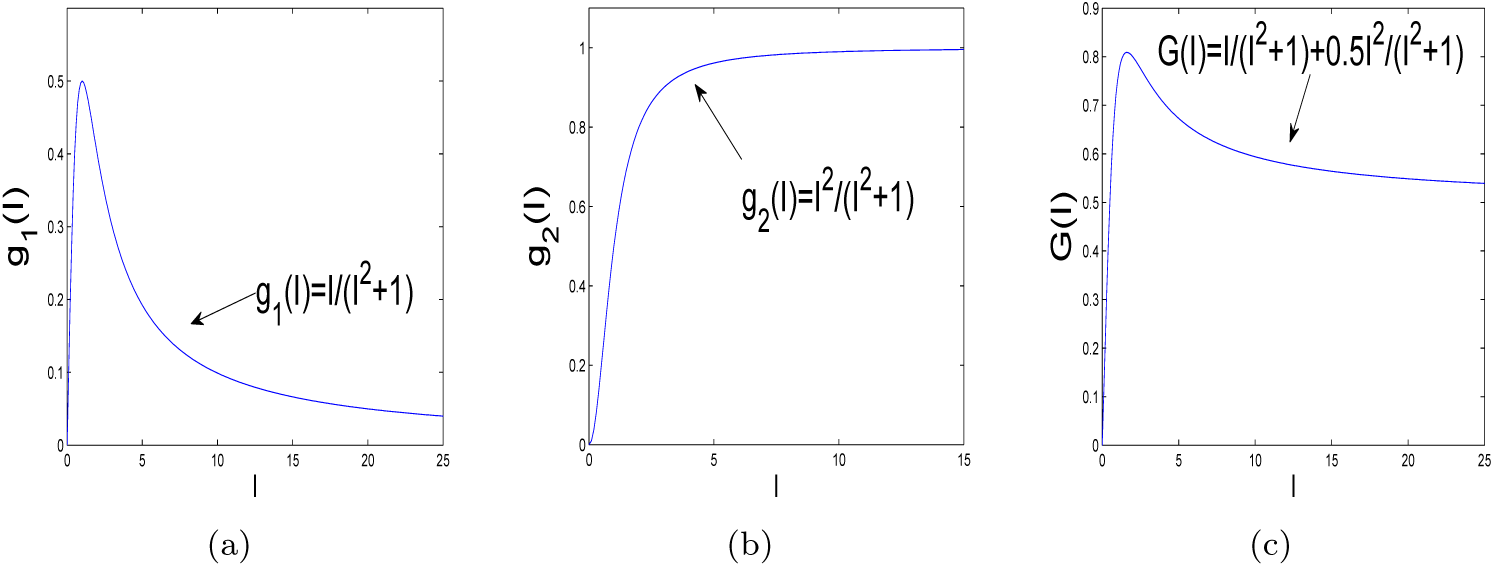
Graphs of different incidence rate functions. (a) Nonmonotone incidence 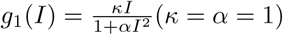. (b) Nonlinear incidence 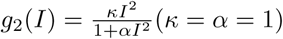. (c) Nonlinear and nonmonotone incidence rate 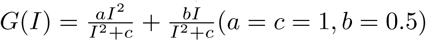.

Furthermore, Li, Zhao and Zhu (see [20]) studied the following SIS model, which describes behavior change effect of susceptible individual when infectious population increases

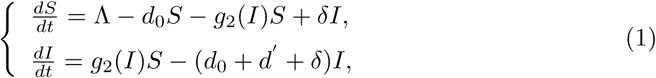

where

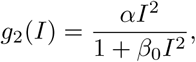

see Fig. 1(b). By the qualitative and bifurcation analyses, they showed that the maximal multiplicity of weak focus is 2, and proved that the model can undergo a Bogdanov-Takens bifurcation of codimension 2. These results illustrate that the behavior change of the susceptible individuals may affect the final spread level of an epidemic.

Actually, both the effect of psychological and behavior affect of susceptible individuals have influence on the transmission of the disease. Thus, motivated by the above researches, we consider a nonlinear incidence rate of a SIS model as follows

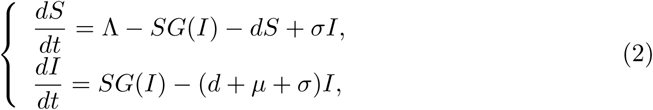

where

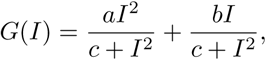

see Fig. 1(c), which can describe the effect of psychology and behavior change of susceptible individuals. *S*(*t*) and *I*(*t*) represent the number of susceptible individuals and infected individuals at time *t*, respectively. Λ is the recruitment rate of population, *d* is the natural death, *μ* is the disease-induced death rate, and *σ* represents the recovered rate, *a, b* and *c* are all positive constants.

The organization of this paper is as follows. First of all, we analyze the existence of equilibria and local stability of equilibria. Then, we study the existence of Hopf bifurcation around the positive equilibrium at the critical value under the conditions of *R*_0_ *<* 1 and *R*_0_ *>* 1. We also show that these positive equilibria can be weak focus for some parameter values and a cusp type of Bogdanov-Takens bifurcation of codimension 3. Finally, we give some brief conclusion.

### Types and Stability of equilibria

In this section, our aim is to perform an elaborative analysis of equilibria for system (2). Firstly, for convenience, we make scalings: 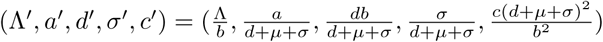, and 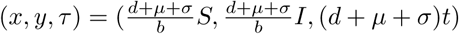. To avoid the abuse of mathematical notation, we still denote (Λ*^′^, a^′^, d^′^, σ^′^, c^′^, τ*) by (Λ, *a, d, σ, c, t*). Then model (2) becomes

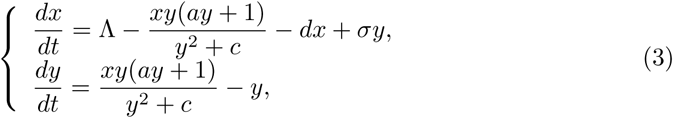

with *d* + *σ <* 1.

#### Lemma 1

*The set 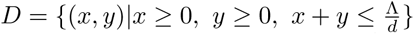 is an invariant manifold of system* (3), *which is attracting in the first octant of* ℝ^2^.

**Proof** Summing up the two equations in (3), we can get

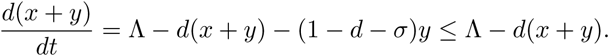

Thus, 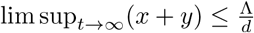, which implies the conclusion.

Obviously, system (3) always has a unique disease-free equilibrium 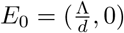. The positive equilibria of (3) can be gained by solving the following algebraic equations

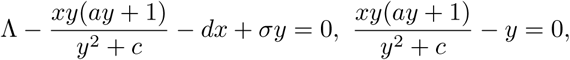

which yields

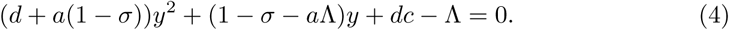

Denote the basic reproduction number as follows,

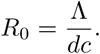

Computing the discriminant of (4), we get

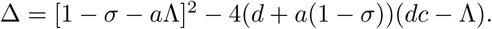

Furthermore, define

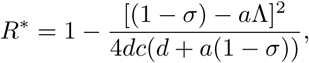

then Δ > 0 if and only if *R*_0_ *> R^∗^*, Δ = 0 if and only if *R*_0_ = *R*^*∗*^ and Δ *<* 0 if and only if *R*_0_ *< R^∗^*. It is obvious that *R*^*∗*^ *<* 1 and we can obtain the following theorem.

#### Theorem 1

*Model* (3) *always has a disease-free equilibrium E*_0_ *and the following conclusions hold.*

*(i) When R*_0_ *<* 1, *we have*

*(a) if R*_0_ *< R^∗^, then system* (3) *has no positive equilibrium;*

*(b) if R*_0_ = *R*^*∗*^ *and* Λ > (1 *- σ*)*/a, then system* (3) *has a unique positive equilibrium*

*E*_*1*_ (*x, y*), *where 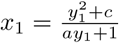 and 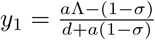;*

*(c)if R*_0_ *> R^∗^ and* Λ > (1 − *σ*)*/a, then system* (3) *has two positive equilibria*

*E*_2_(*x*_2_, *y*_2_) *and E*_3_(*x*_3_, *y*_3_), *where* 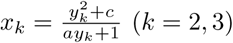 (*k* = 2, 3) *and 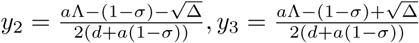;*

*(ii)When R*_0_ = 1 *and* Λ > (1 *- σ*)*/a, then system* (3) *has a unique positive equilibrium E_5_*(*x*_*5*_, *y*_*5*_), *where 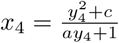 and 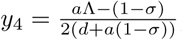.*

*(iii)When R*_0_ > 1, *then system* (3) *has a unique positive equilibrium E*_5_(*x*_5_, *y*_5_), *where* 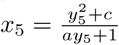 and 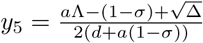.

In the following, we discuss the local stability of *E*_*k*_(*x*_*k*_, *y*_*k*_) (*k* = 0, 1, 2, 3, 4, 5) and present the corresponding phase portrait. By directly calculating, the Jacobian matrix at equilibrium *E*_*k*_ is

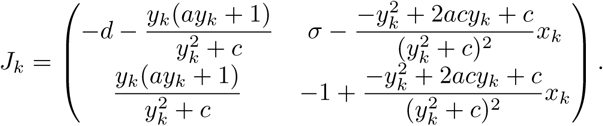

We have

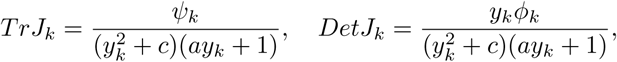

where

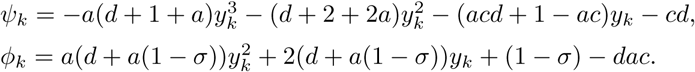

In addition, after some complicated computations, we get

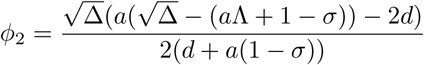

and

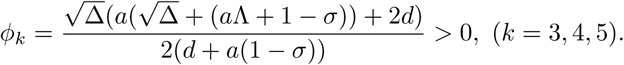

#### Theorem 2

*The disease free equilibrium E*_0_ *of system* (3) *is*

*(i) an attracting node if R*_0_ *<* 1;

*(ii) a hyperbolic saddle if R*_0_ > 1;

*(iii) a saddle-node of codimension 1 if R*_0_ = 1 *and* 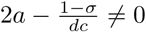; *a repelling node if R*_0_ = 1 *and* 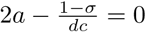.

**Proof** Obviously, at equilibrium *E*_0_, we have det(*J*_0_) = *d*(1 *- R*_0_) and Tr(*J*_0_) = *-d -* (1 *- R*_0_). Therefore, *E*_0_ is a stable node if *R*_0_ *<* 1 and a hyperbolic saddle if *R*_0_ > 1, and degenerate if *R*_0_ = 1.

When *R*_0_ = 1, we let 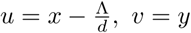, *v* = *y*, then system (3) becomes

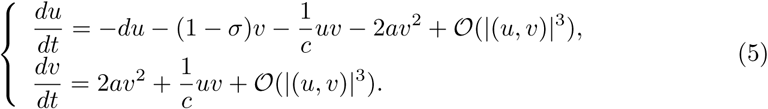

Indeed, if *R*_0_ = 1, the Jacobian *J*_0_ is diagonalizable with eigenvalues *λ*_1_ = *-d* and *λ*_2_ = 0 and respective eigenvectors *ν*_1_ = (1, 0) and 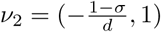. By the transformation

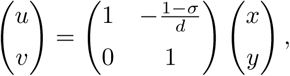

system (5) can be rewritten as

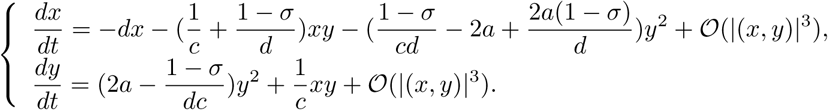

If 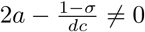, according to the calculation of center manifold [15], we know that the center manifold *x* = *h*(*y*) of (5) begins with quadratic term of *y*. In addition, from the second equation of (5), we can easily obtain that the equation restricted to the center manifold as follows

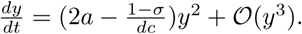

By applying the Theorem 7.1 in Zhang et al. [15], *E*_0_ is a saddle-node.

If 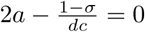, then the center manifold turns into

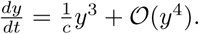

Since 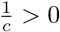 and the first nonzero item is uneven. Thus, the equilibrium *E*_0_ is a repelling node, according to Theorem 7.1 in Zhang et al. [15].

From the expression of *ψ*_1_ and *ϕ*_1_, we can see that one of the eigenvalues of characteristic matrix of *E*_1_ is zero and the other is nonzero if *ψ*_1_ = 0. The type of *E*_1_ can be directed proved by checking the conditions in Zhang et al. ([15], Theorem 7.1-7.3). So, we have the following results.

#### Theorem 3

*If R*_0_ = *R*^*∗*^, *then system* (3) *has a unique positive equilibrium E*_1_. *More Precisely,*

*(a) if ψ*_1_ *≠* 0, *then E*_1_ *is a saddle-node;*

*(b) if ψ*_1_ = 0, *then E*_1_ *is a cusp.*

#### Theorem 4

*Suppose that R^∗^ < R*_0_ *<* 1 *and a*Λ > 1 *- σ, then system* (3) *has two positive equilibria E*_2_ *and E*_3_, *and equilibrium E*_2_ *is a hyperbolic saddle for all permissible choices of parameters, equilibrium E*_3_ *is not degenerate. Moreover,*

*(i) E*_3_ *is a stable focus or node if ψ*_3_ *<* 0;

*(ii) E*_3_ *is a weak focus or center if ψ*_3_ = 0;

*(iii) E*_3_ *is an unstable focus or node if ψ*_3_ > 0.

**Proof** Note that *ϕ*_2_ is less than zero since Δ *<* (*a*Λ + 1 *- σ*)^2^, then *E*_2_ is a hyperbolic saddle for any choices of parameters. And at *E*_3_, we have *ϕ*_3_ > 0. Thus, the stability of equilibrium *E*_3_ depends on the sign of *ψ*_3_.

#### Theorem 5

*When R*_0_ = 1 *and 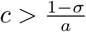, then system* (3) *has a unique positive equilibrium E*_4_(*x*_4_, *y*_4_), *and the equilibrium E*_4_ *is stable if ψ*_4_ *<* 0.

**Proof** In fact, when *R*_0_ = 1 and 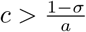, then *ϕ*_4_ > 0. Thus, the stability of *E*_4_ is determined by the sign of *ψ*_4_.

#### Theorem 6

*Assume R*_0_ > 1, *then system* (3) *has a unique positive equilibrium E*_5_. *Moreover,*

*(i) E*_5_ *is stable if ψ*_5_ *<* 0;

*(ii) E*_5_ *is a weak focus or center if ψ*_5_ = 0;

*(iii) E*_5_ *is unstable if ψ*_5_ > 0.

**Proof** Obviously, when *R*_0_ > 1, then *ϕ*_5_ > 0, and then *E*_5_ is stable if *ψ*_5_ *<* 0.

#### Lemma 2

*From the expression of ψ_k_* (*k* = 1, 3, 5), *we can obtain that E_k_* (*k* = 1, 3, 5) *is always stable if 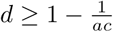.*

When *ψ*_*i*_ 0 (*i* = 3, 5), the dynamics of system (3) can be easily seen in Fig. 2, Fig. 3 and Fig. 4, respectively. The dynamical behaviors of system (3) when *ψ*_*i*_ = 0 (*i* = 3, 5) will be discussed in detail in the next section.

**Fig 2:**
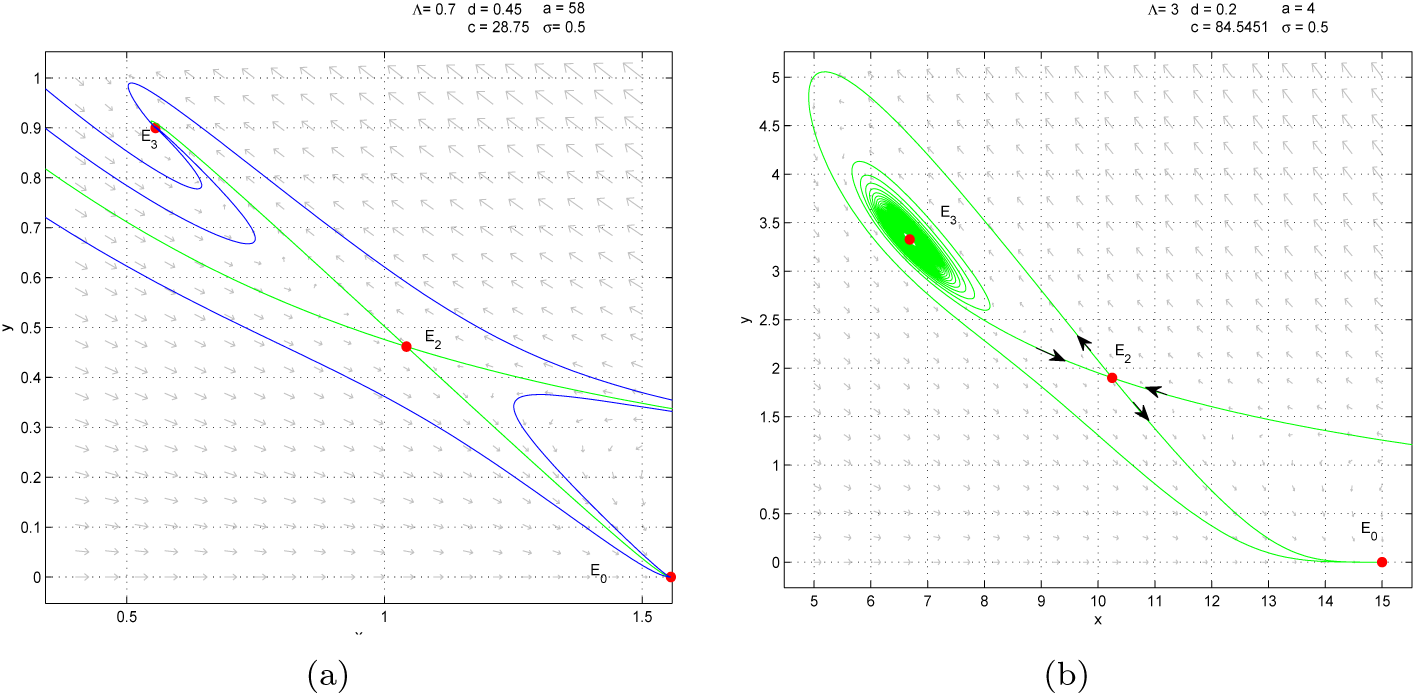
**When** *R*^*∗*^ *< R*_0_ *<* 1, Λ > (1 *- σ*)*/a*. (*a*): *ψ*_3_ *<* 0, equilibria *E*_0_ and *E*_3_ are locally stable and *E*_2_ is unstable. (*b*): *ψ*_3_ > 0, equilibrium *E*_0_ is locally stable and *E*_2_ and *E*_3_ are unstable.

**Fig 3:**
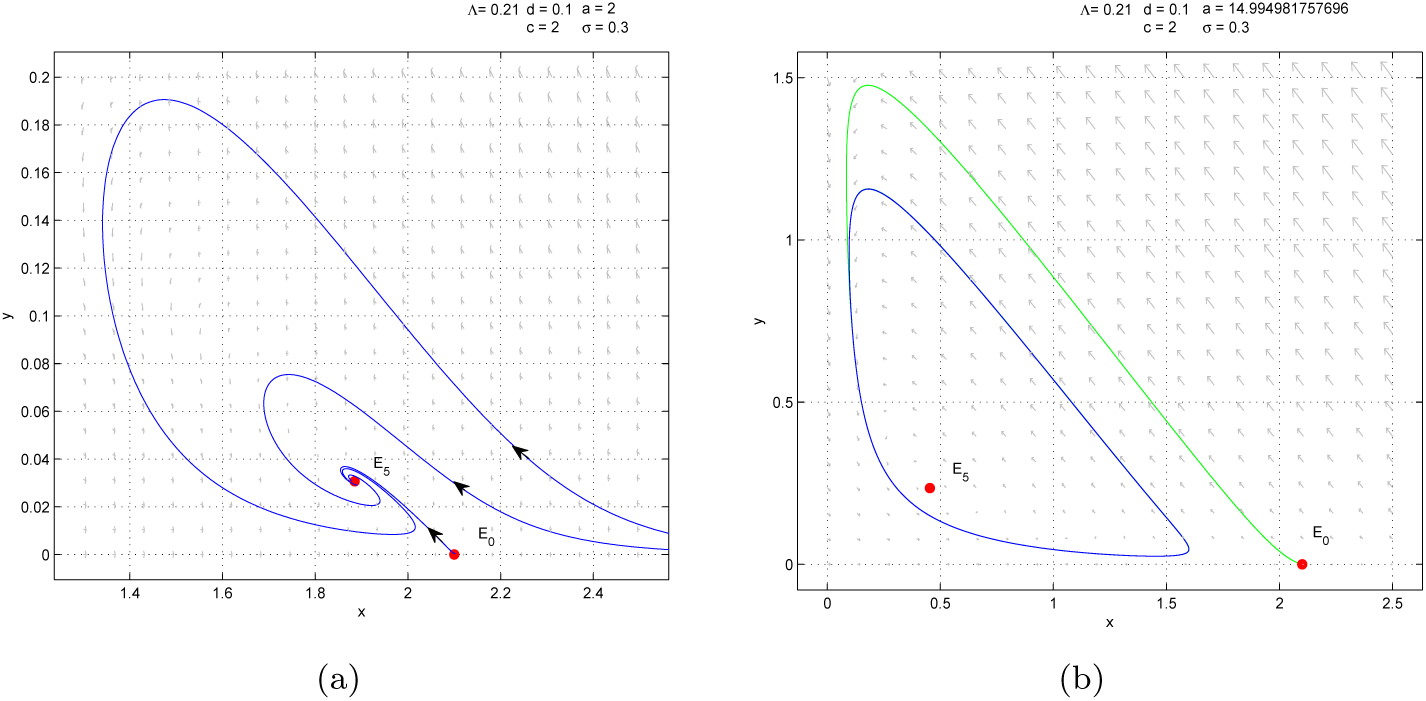
**When** *R*_0_ > 1. (*a*): *ψ*_5_ *<* 0, the disease-free equilibrium *E*_0_ is unstable and the unique positive equilibrium *E*_5_ is globally stable. (*b*): *ψ*_5_ > 0, *E*_0_ and *E*_5_ are unstable and there exists a stable limit cycle.

**Fig 4:**
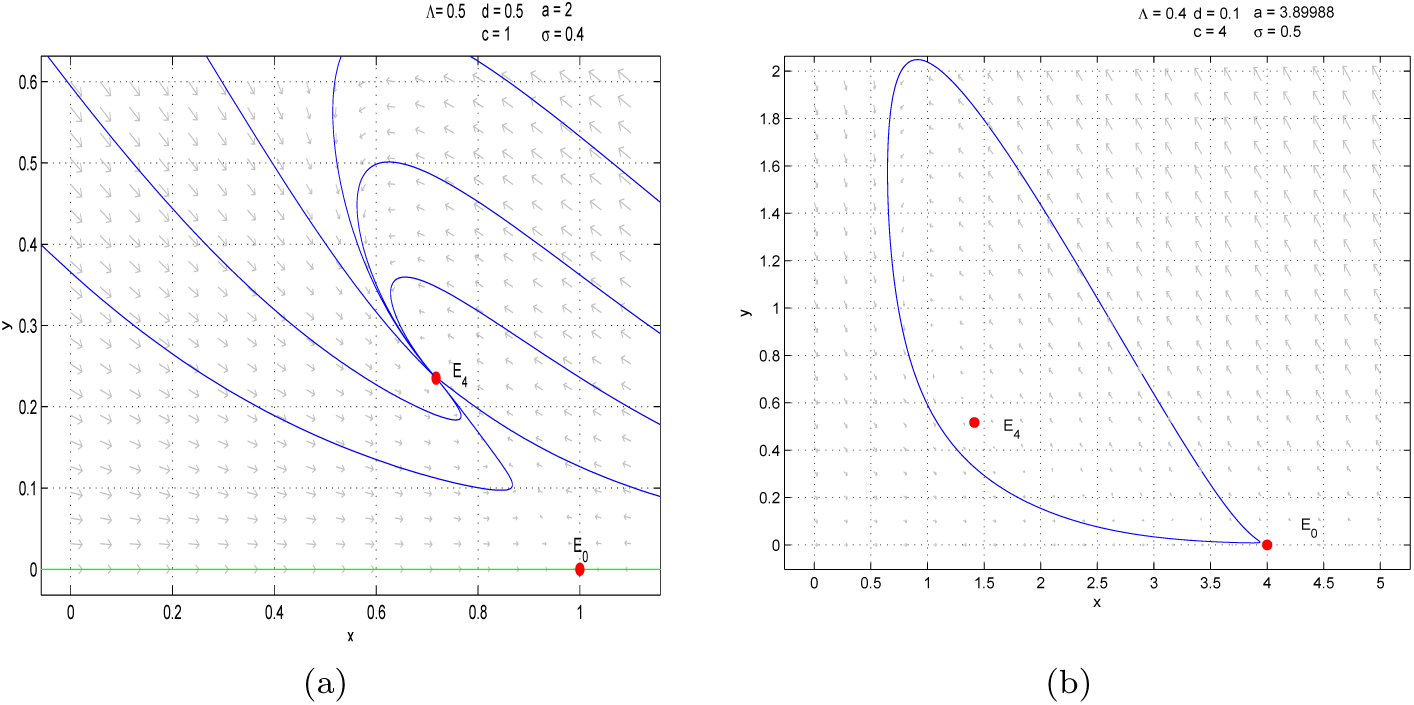
**When** *R*_0_ = 1**, and** Λ > (1 *- σ*)*/a*. (*a*): *ψ*_4_ *<* 0, the disease-free equilibrium *E*_0_ is unstable and the unique positive equilibrium *E*_4_ is globally stable. (*b*): *ψ*_4_ > 0, *E*_0_ and *E*_4_ are unstable and there exists a stable limit cycle.

**Remark 1** *In fact, Fig. 2 (a) shows the occurrence of bi-stability, in which solution may converge to one of the two equilibria, depending on the initial conditions. And in practical, this interesting phenomenon implies that initial states determine whether the disease can dies out or not.*

**Remark 2** *From Fig. 2 (a), we can see there exists two seperatrices. All solutions tend to the disease-free equilibrium E*_0_ *except the two green lines tend to equilibrium E*_2_.

#### Theorem 7

*Suppose that R*_0_ = 1 *and* 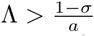, *then system* (3) *has a unique positive equilibrium E*_4_. *If ψ*_4_ > 0, *then there exists at least one stable limit cycle in the interior of the first quadrant.*

**Proof** Indeed, the Jacobian *J*_0_ is diagonalizable with eigenvalues *λ*_1_ = *-d* and *λ*_2_ = 0 and respective eigenvectors *ν*_1_ = (1, 0) and 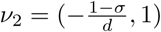. By the transformation

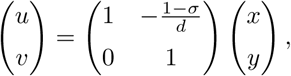

system (5) becomes

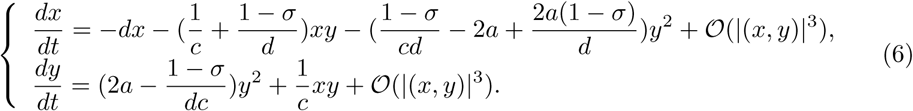

The theorem of Chochitaichvili [21] yields directly that system (6) is topologically equivalent to the system:

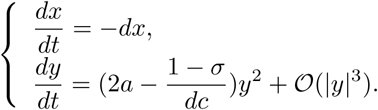

It is easy to see that 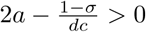 according to the condition of *R*_0_ = 1 and 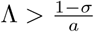, and then we can gain that there exist a unique repelling equilibrium *E*_4_ in the region *D*_1_ shown in (*a*) of Fig.5. Consequently, by the Poincar*é*-Bendixson theorem, at least one stable limit cycle appears in the interior of the first quadrant.

**Fig 5:**
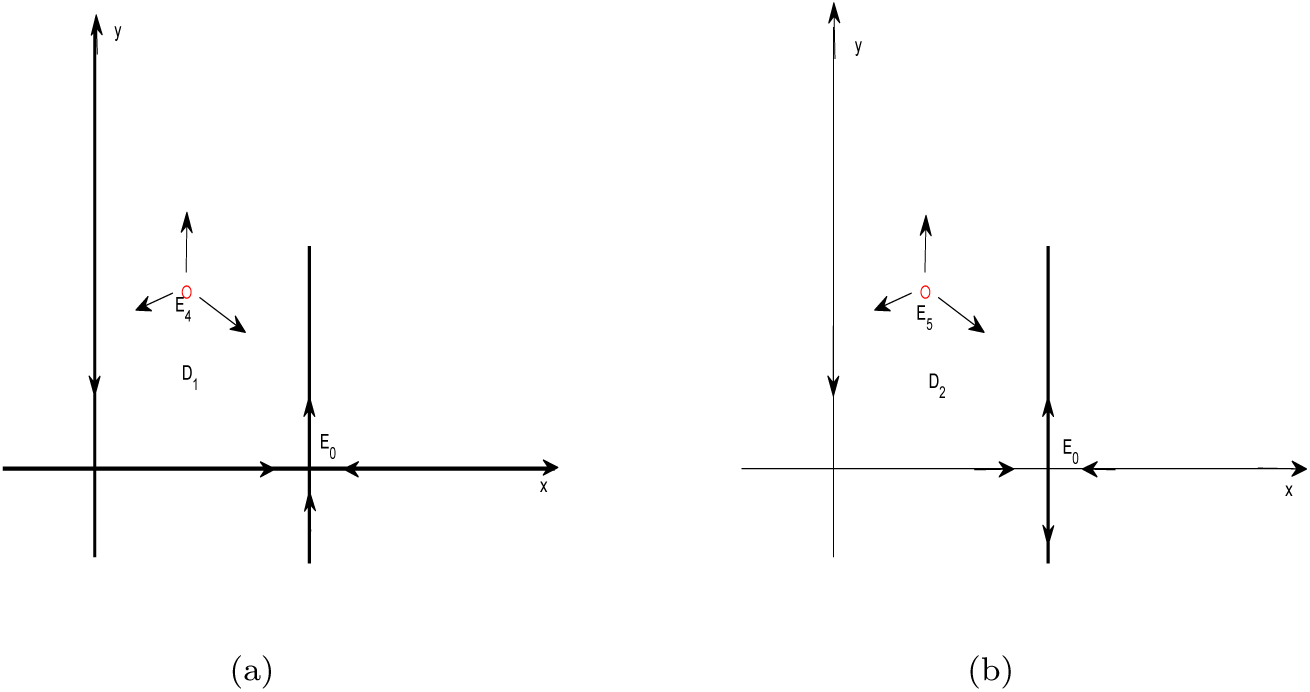
**When** *R*_0_ = 1**, and** λ > (1 *- σ*)*/a*. (*a*): When *R*_0_ = 1, 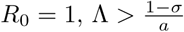 and *ψ*_4_ > 0, there exists a stable limit cycle. (*b*): When *R*_0_ > 1 and *ψ*_5_ > 0, there exists a stable limit cycle.

Similarly, when *ψ*_5_ > 0, we have the following result.

#### Theorem 8

*Suppose that R*_0_ > 1. *If ψ*_5_ > 0, *then there exists at least one stable limit cycle in the interior of the first quadrant.*

**Proof** Indeed, the Jacobian *J*_5_ has eigenvalues *λ*_1_ = *-d* and *λ*_2_ = *R*_0_ *-* 1 > 0, when *R*_0_ > 1. Thus, we can gain that there exist a *E*_5_, which is the unique repelling equilibrium in the region *D*_2_ shown in (*b*) of Fig.5. Consequently, by the Poincar*é*-Bendixson theorem, at least one stable limit cycle appears in the interior of the first quadrant.

**Remark 3** *From* (*b*) *of Fig. 3, we can see clearly that there exist a stable limit cycle enclosing the equilibrium E*_5_ *even though E*_0_ *is a saddle-node. Similarly, from* (*b*) *of Fig. 4, we can also gain that there exists a stable limit cycle if E*_4_ *is unstable.*

### Bifurcations Backward bifurcation

#### Theorem 9

*When R*_0_ = 1 *and* 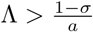, *model* (3) *exhibits a backward bifurcation at equilibrium E*_0_.

**Remark 4** *When R*_0_ = 1 *and* 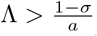, *system* (3) *exhibits a unique positive equilibrium E*_4_, *which means that once R*_0_ *crosses 1, the disease can invade to a relatively high level. And this is one of the main characters of backward bifurcation [19].*

**Remark 5** *Actually, backward bifurcation did not emerge with a* = 0, *which is considered in [20]. This indicates that introducing the non-motonic incidence into model makes the epidemic model more complex and exhibit richer dynamical behaviors.*

### Hopf bifurcation

In this subsection, we will study the Hopf bifurcation of system (3) for (i) *R*^*∗*^ *< R*_0_ *<* 1 and λ > (1 *- σ*)*/a*; (ii) *R*_0_ > 1. From the discussion in Section 2, it can be seen that Hopf bifurcation may occur at *E*_3_, *E*_5_. As the expressions of equilibria *E*_3_ and *E*_5_ are same, no considering the values of every parameters. Based on Theorem 4, Theorem 5 and Theorem 6, we know that the stability of *E*_3_ and *E*_5_ are similar and when *ψ*_*k*_ = 0, *E*_*k*_(*k* = 3, 5) is a weak focus or center. Thus, we show the existence of Hopf bifurcation around *E*_*k*_(*k* = 3, 5).

#### Theorem 10

*Suppose E_k_* (*k* = 3, 5) *exist, then model* (2) *undergoes a Hopf bifurcation at equilibrium E_k_ if ψ_k_* = 0. *Moreover,*

*(a) if η <* 0, *there is a family of stable periodic orbits of model* (3) *as ψ_k_ decreases from* 0;

*(b) if η* = 0, *there are at least two limit cycles in* (3), *where η will be defined below;*

*(c) if η >* 0, *there is a family of unstable periodic orbits of* (3) *as ψ_k_ increases from* 0.

**Proof** From the above discussions, we can see that *trJ*_*k*_ = 0 if and only if *ψ*_*k*_ = 0, and det *J*_*k*_ > 0 when equilibrium *E*_*k*_ exists. Therefore, the eigenvalues of *J*_*k*_ are a pair of pure imaginary roots if *ψ*_*k*_ = 0. From direct calculations we have that

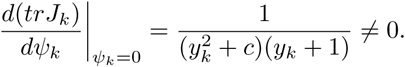

By Theorem 3.4.2 in [16], ψ*_k_* = 0 is the Hopf bifurcation point for (3).

Next, introduce a new time variable *τ* by *dt* = (*y*^2^ + *c*)*dτ*. By rewriting *τ* as *t*, we obtain the following equivalent system of (3)

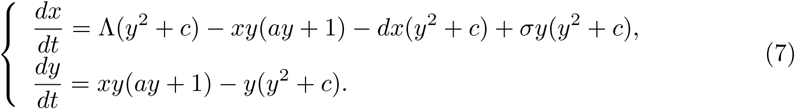

Let *X* = *x - x_k_* and *Y* = *y - y_k_*, still use (*x, y*) to express (*X, Y*), then system (7) becomes

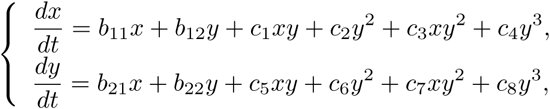

where

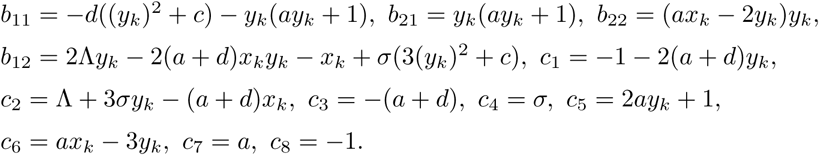

Let 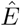 denote the origin of *x - y* plane. Since *E*_*k*_ satisfies Eq. (3), we obtain

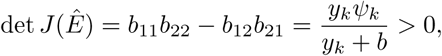

and it is easy to verify that *b*_*11*_ + *b*_*22*_ = *0* if and only if *ψ*_*k*_ = 0. Let 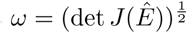, *u* = *-x* and 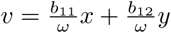, then the normal form of system (7) reads

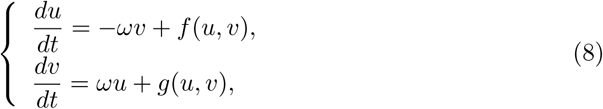

where

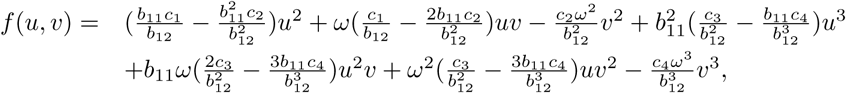

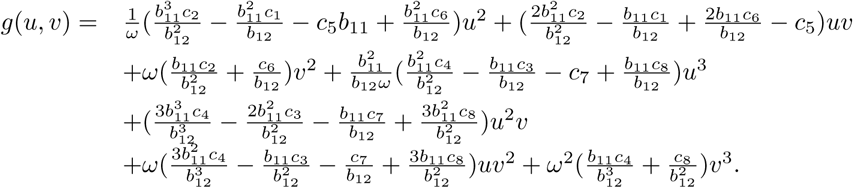

Set

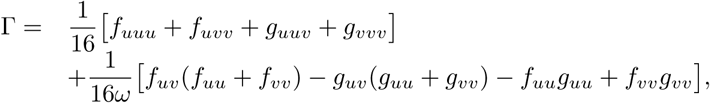

where *f*_*uv*_ denotes 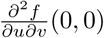, etc. Then, by computing we obtain

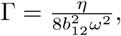

where

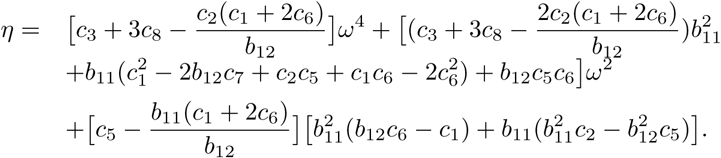

By Theorem 3.4.2 and Theorem 3.4.11 in [16], the rest claims in Theorem 10 are true.

**Remark 6** *What need to be noted here is that the expression b*_12_ *in Theorem 10 is nonzero. Otherwise, we have* det *J*_*k*_ = *b*_11_*b*_22_ *<* 0, *since b*_11_ + *b*_22_ = 0, *which is an contradiction.*

Next, we present examples to show that equilibrium *E*_*k*_ can be a stable weak focus of multiplicity two, and under a small perturbation, system (3) undergoes a degenerate Hopf bifurcation and produces two limit cycles.

Firstly, fix *y*_*k*_ = 1/2 and solve for 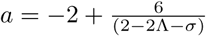. Also, fix *x*_*k*_ = 1, based on 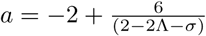, we can get 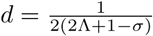.Then Ψ*_k_* = 0 if and only if 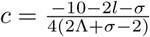. Secondly, substitute these expressions into *η* and through complicated computation, we can obtain

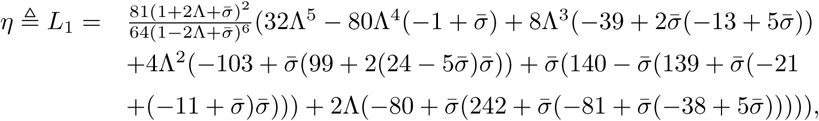

with 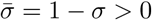, which is the first Liapunov number of the equilibrium (0, 0) of (8). Then, we fix 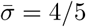 and solve equation *η* = 0, then we get only one suitable value 0.5859 for λ. That is to say, if (*σ,* λ, *d, a, c*) = (1/5, 5859, 0.2, 8, 4.741), then *L*_1_ = 0. Furthermore, it can be seen that *E*_*k*_ = *E*_3_ under this group of parameters.

In the following, we further compute the second Liapunov number of the equilibrium (0, 0) of system (8) by the successor function method. It is convenient to introduce polar coordinates (*r, θ*) and rewrite system (8) in polar coordinates by *x* = *r* cos *θ, y* = *r* sin *θ*. It is clear that in a small neighborhood of the origin, the successor function *D*(*c*_0_) of system (8) can be expressed by

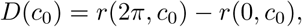

where *r*(*θ, c*_0_) is the solution of the following Cauchy problem

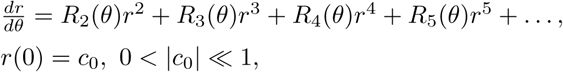

where *R*_*i*_(*θ*) (*i* = 1, 2, 3 *…*) is a polynomial of (sin *θ,* cos *θ*), whose coefficient can be expressed by the coefficients of system (8). We omit them here, since the expressions are too long.

With the aid of Mathematica, we get

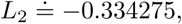

and

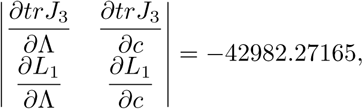

when (*σ,* Λ, *d, a, c*) = (1/5, 5859, 0.2, 8, 4.741). Therefore, the interior equilibrium *E*_3_ is a stable weak focus of multiplicity two if (*σ,* λ, *d, a, c*) = (1/5, 5859, 0.2, 8, 4.741). The phase portraits of system (3) under this group of parameters are shown in Fig. 6.

**Fig 6:**
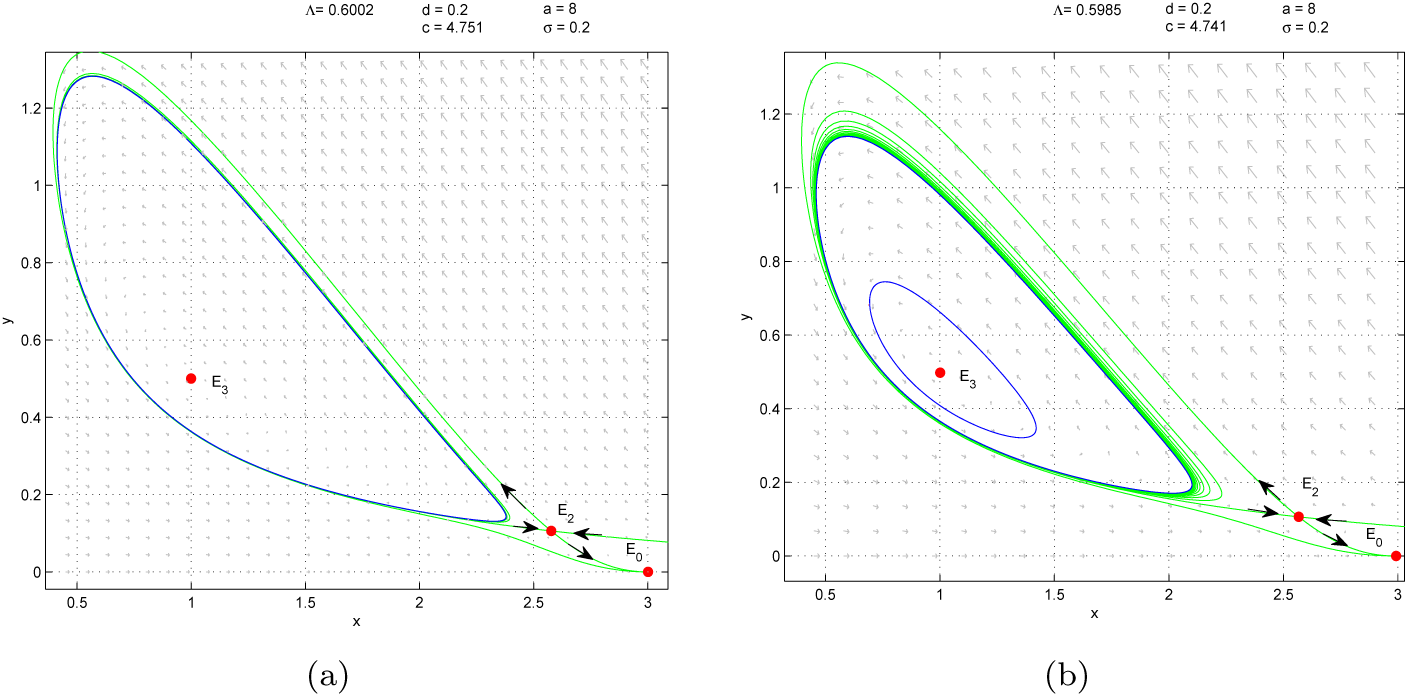
**Bold the figure title.** (*a*): One Limit cycles enclosing an unstable hyperbolic focus *E*_3_. (*b*): Two Limit cycles enclosing an unstable hyperbolic focus *E*_3_, and the small one is stable, the large one is unstable.

Besides, we give the numerical simulation graphs for one limit cycle and two limit cycles under small perturbations of some parameters. From Fig. 6 (*a*), we can see that there exist only one limit cycle around the endemic *E*_3_, and Fig. 6 (*b*) shows us that a new limit cycle emerge with small perturbations of parameters λ and *c*. It is worth emphasizing that if we change the values of parameters λ and *c*, an unstable homoclinic loop arise, which is shown in Fig.7.

**Fig 7:**
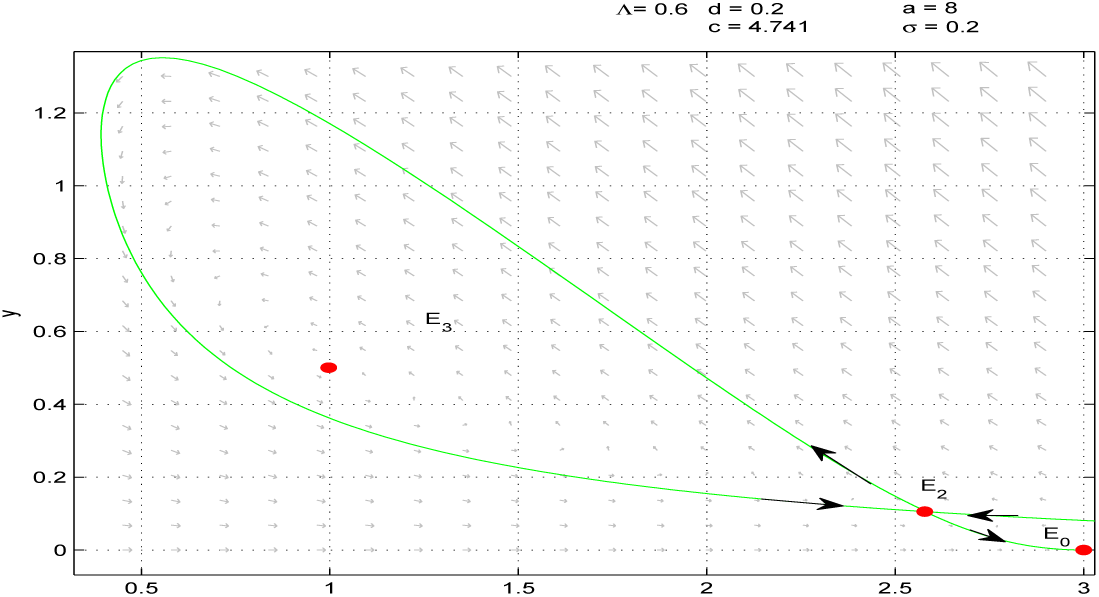
Equilibrium *E*_3_ is stable and there exists an unstable homoclinic loop.

Around equilibrium *E*_5_, we can gain the same result from Fig. 8. Under the condition that parameter *a, d, σ* take value 5.6923, 0.11, 0.2 and change value of λ and *c* from 0.45102, 3.6062 to 0.51, 3.5972, respectively, which is a minor change, then the number of limit cycles will add one. Thus, the interior equilibrium *E*_5_ is a stable weak focus of multiplicity two if (*σ,* λ, *d, a, c*) = (0.2, 0.51, 0.11, 5.6923, 3.5962).

**Fig 8:**
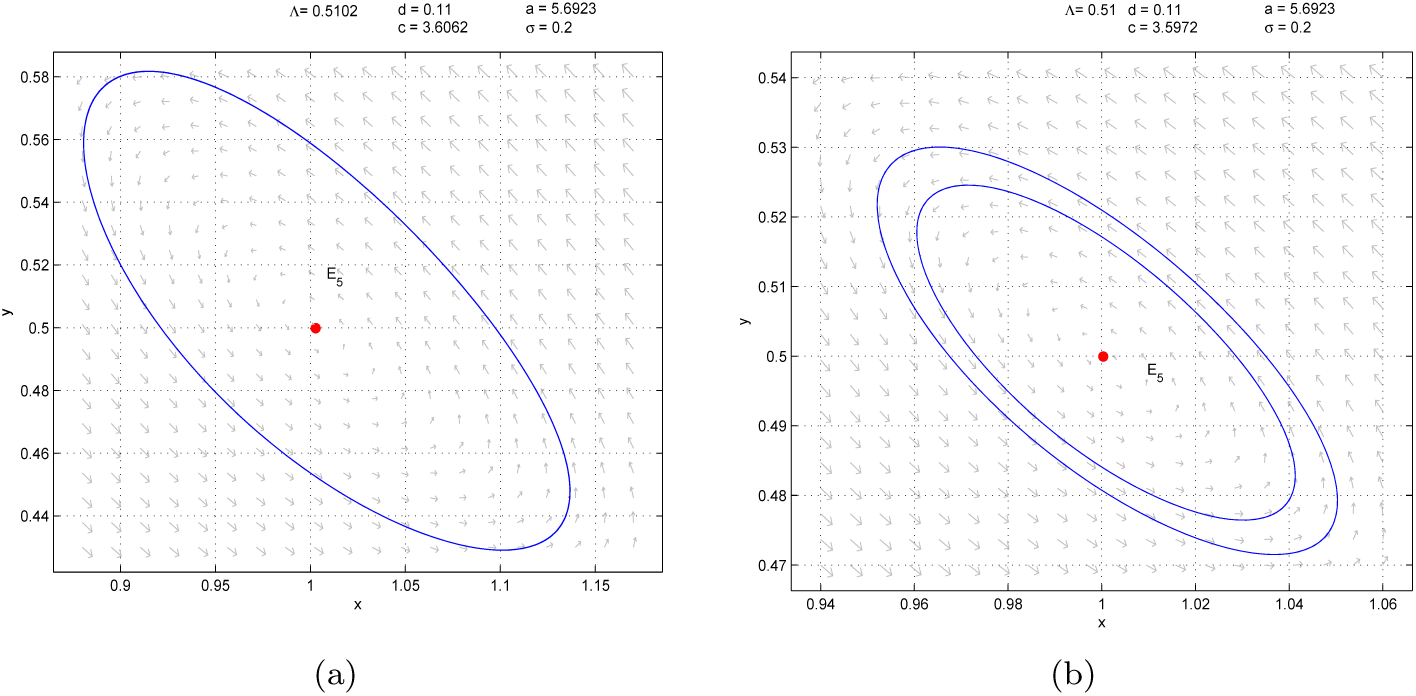
(*a*): One Limit cycles enclosing an unstable hyperbolic focus *E*_5_. (*b*): Two Limit cycles enclosing an unstable hyperbolic focus *E*_5_, and the small one is stable, the large one is unstable.

**Remark 7** *As a matter of fact, the reproduction number is equal to zero in [8, 20], which simplifies the condition that Hopf bifurcation occur. In our model, we also comprehensively discuss the existence of Hopf bifurcation when R*_0_ *<* 1, *R*_0_ = 1 *and R*_0_ > 1. *Besides, authors in [20] did not show the appearance of homoclinic loop, which is an interesting bifurcation phenomenon given in Fig.7.*

### Bogdanov-Takens bifurcation

In this subsection, we investigate the Bogdanov-Takens bifurcation in system (3). The following Lemma 3 is from Perko [17], and Lemma 4 is Proposition 5.3 in Lamontage et al. [18].

#### Lemma 3

*The system*

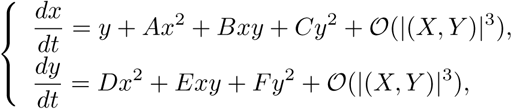

*is equivalent to the system*

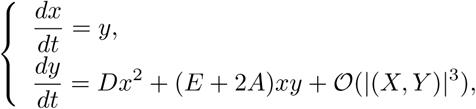

*in some small neighborhood of* (0, 0) *after changes of coordinates.*

#### Lemma 4

*The system*

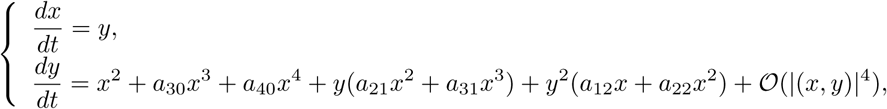

*is equivalent to the system*

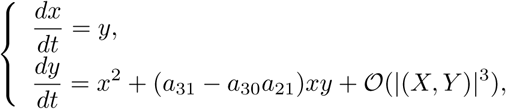

*in some small neighborhood of* (0, 0) *after changes of coordinates.*

From Theorem 1, we can see that there exists a unique positive equilibrium *E*_1_(*x*_1_, *y*_1_) when *R*_0_ = *R*^*∗*^, where

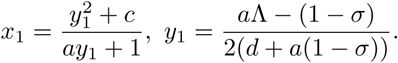

From the proving process of Theorem 4, it is easily to know that *detJ*_1_ = 0. And Theorem 3 suggests that the characteristic matrix *J*_1_ possesses a zero eigenvalue with multiplicity 2 when *ψ*_1_ = 0, which shows that system (3) may admit a Bogdanov-Takens bifurcation. Thus, we can prove the following theorem.

Define two functions:

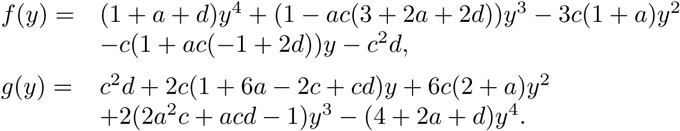

#### Theorem 11

*Suppose that R*_0_ = *R*^*∗*^ and *ψ*_1_ = 0, *then the only interior equilibrium E*_1_ *of system* (3) *is a cusp. Moreover,*

*(a) if f* (*y*_1_)*g*(*y*_1_) *≠* 0, *then E*_1_ *is a cusp of codimension 2;*

*(b) if f* (*y*_1_)*g*(*y*_1_) = 0, *then E*_1_ *is a cusp of codimension greater than or equal to 3.*

**Proof** Changing the variables as *X* = *x - x*_1_, *Y* = *y - y*_1_, system (3) becomes

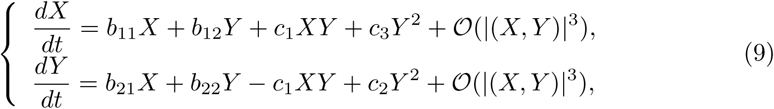

where

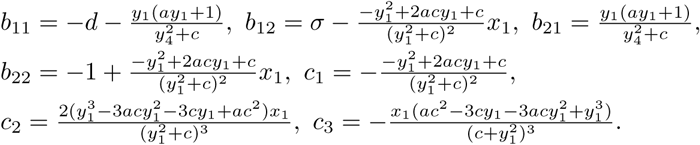

Make the non-singular linear transformation

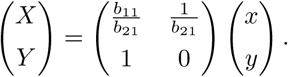

Then system (9) is transformed into

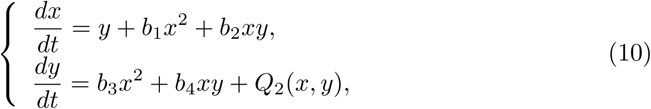

where *Q*_2_(*x, y*) is a smooth function in (*x, y*) at least of the third order and

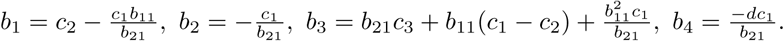

By Lemma 3, we obtain an topologically equivalent system of system (10)

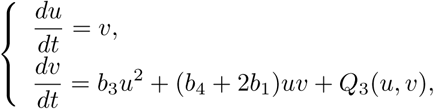

where

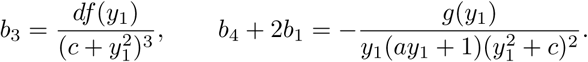

Therefore, *E*_1_ is a cusp of codimension 2 if *f* (*y*_1_)*g*(*y*_1_) ≠ 0, by the results in Perko [17], or else, *E*_1_ is a cusp of codimension at least 3.

**Remark 8** *In fact, Theorem 11* (*b*) *include the following three cases:*

(1) *if f* (*y*_1_) 0 *and g*(*y*_1_) = 0, *E*_1_ *is a cusp point;*

(2) *if f* (*y*_1_) = 0 *and g*(*y*_1_) *≠* 0, *E*_1_ *is nilpotent focus/elliptic point;*

(3)*if f* (*y*_1_) = *g*(*y*_1_) = 0, *E*_1_ *is a nilpotent focus.*

Unfortunately, because of the complexity of *f* (*y*_1_) and *g*(*y*_1_), we can not determine which of these three situations can occur theoretically. But we will show for some parameter values, *f* (*y*_1_) = 0 and *g*(*y*_1_) = 0, *i.e. E*_1_ is a cusp point.

In the following, we will give an example to show that the case (*b*) of Theorem 11 occur.

In the first place, fix *y*_1_ = 1/5, then from 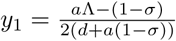, we can solve for 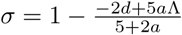. By assumptions *ψ*_1_ = 0 and *R*_0_ = *R*^*∗*^, then from

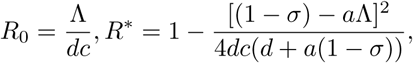

we can get *d* and Λ, where

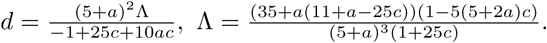

Then

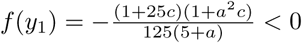

for any *a, c >* 0. We solve *g*(*y*_1_) = 0 for parameter *c* denoted by 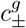, where

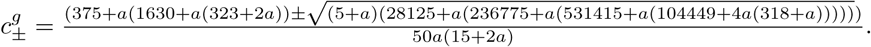

Actually, if 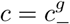, we have *d* = *d*(*a*) *<* 0 for any *a >* 0. Thus, *g*(*y*_1_) = 0 if and only if 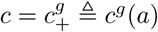. In addition, with the help of *mathematica*, for any 0 *< a <* 1.86433 or *a >* 123.449, one can have all other parameters positive and satisfing *d* + *σ <* 1.

For example, take *a* = 1, then we can get

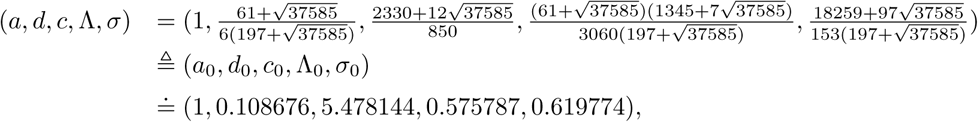

which satisfies *c* = *c*^*g*^(*a*), *i.e.* Theorem 11 (*b*) occur.

#### Theorem 12

*When* (*a, d, c,* λ, *σ*) = (*a*_0_, *d*_0_, *c*_0_, λ_0_, *σ*_0_), *then E*_1_ *is a Bogdanov-Takens point of codimension 3, and the system* (3) *localized at E*_1_ *is topologically equivalent to*

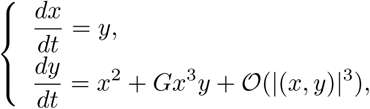

*where G <* 0.

**Proof** First of all, applying a linear transformation *T*_1_: (*x, y*) *→* (*u, v*), defined by *u* = *x - x*_1_, *v* = *y - y*_1_, we can reduce system (3) further to the form

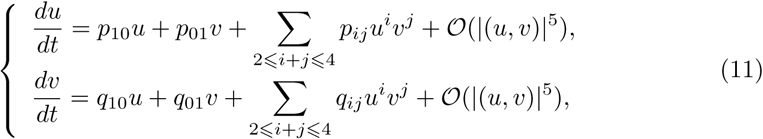

where

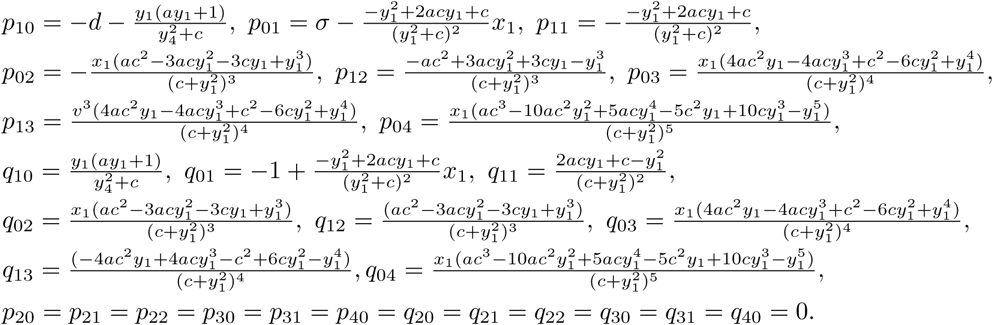

Another transformation *T*_2_: (*u, v*) *→*(*x, y*), defined by *x* = *v, y* = *q*_10_*u - p*_10_*v*, reduces system (11) to

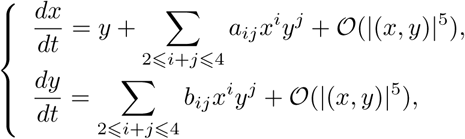

where

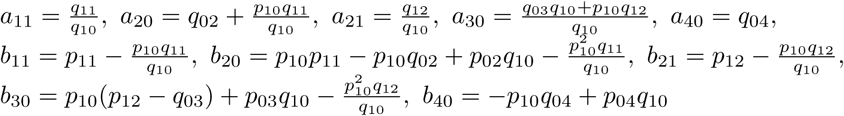

and other coefficients are equal to zero.

Then using the near-identity transformation *T*_3_: (*x, y*) *→* (*u, v*), defined by 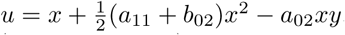, and parameters (*a*_0_, *d*_0_, *c*_0_, λ_0_, *σ*_0_) make *b*_11_ + 2*a*_20_ = 0, so we obtain

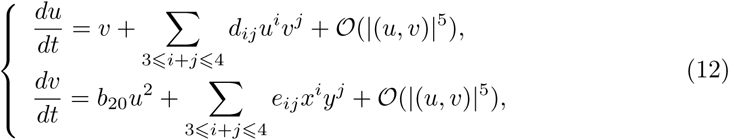

where

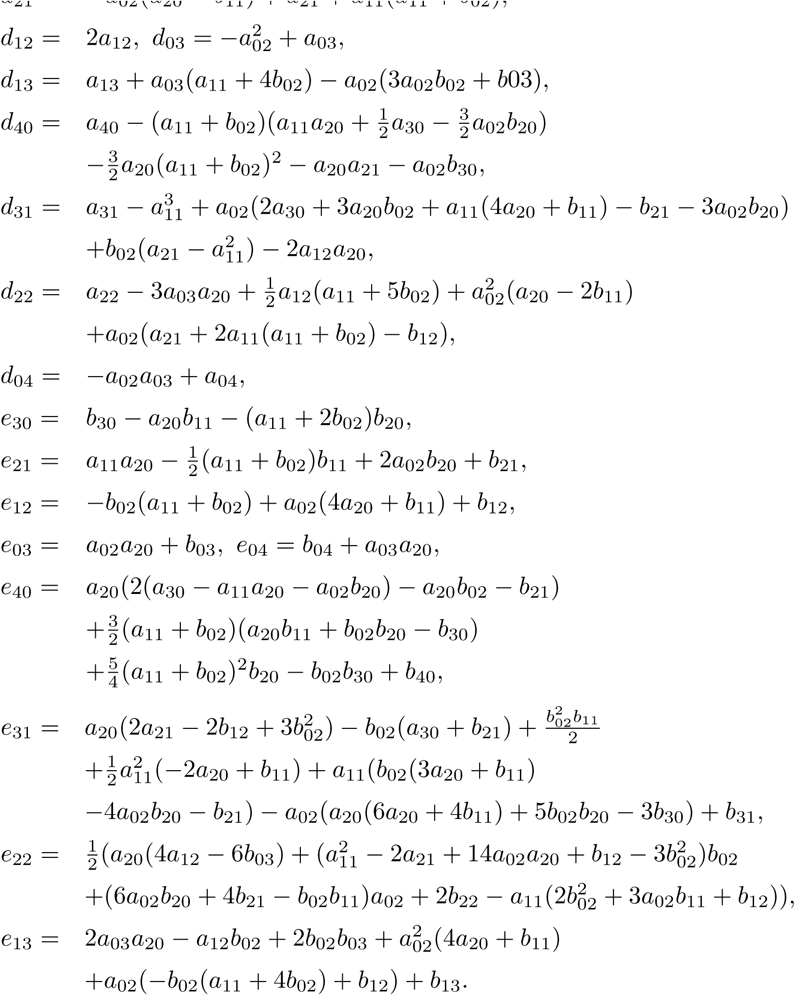

We perform a near-identity smooth change of coordinates

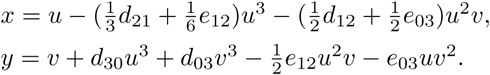

Then system (12) becomes

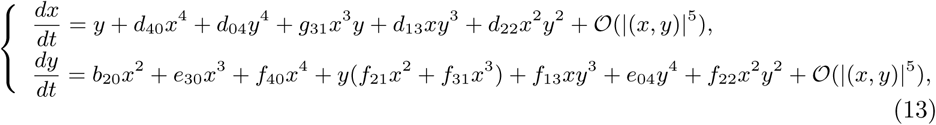

where

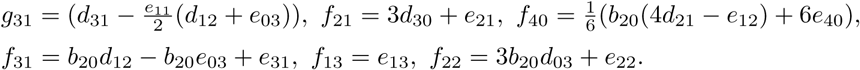

In order to kill the non-resonant cubic terms of system (13), we let

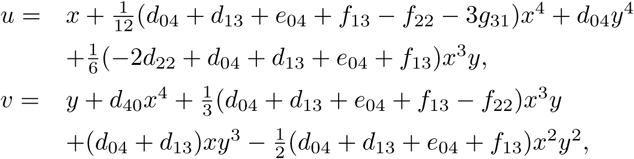

system (13) becomes

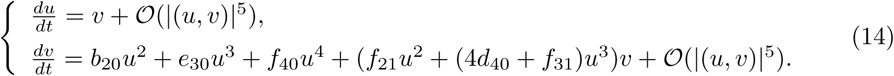

Finally, we set *x* = *b*_20_*u, y* = *b*_20_(*v* + *𝒪*(*|*(*u, v*)*|*5)), system (14) becomes

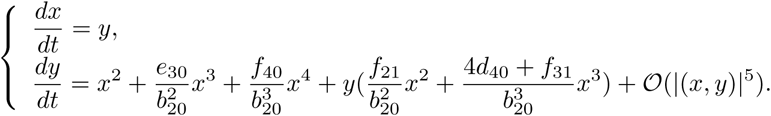

According to Lemma 4 the above system is equivalent to the system

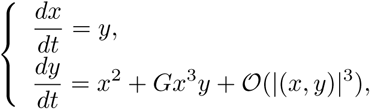

where 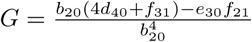. Compute coefficients *b*_20_, *d*_40_, *f*_31_, *e*_30_, *f*_21_ with the condition (*a, d, c,* λ, *σ*) = (*a*_0_, *d*_0_, *c*_0_, λ_0_, *σ*_0_), and straightforward calculation leads to *G* = *-*1.061119*×* 10^6^ *<* 0. Thus, *E*_1_ is a cusp type of Bogdanov-Takens singular with codimension 3.

**Remark 9** *Authors in [8, 20] proved their epidemic model with saturated incidence rate, undergo a Bogdanov-Takens bifurcation of codimension 2. When we consider the incidence of a combination of the saturated incidence rate and a non-monotonic incidence, the codimension of Bogdanov-Takens bifurcation can be up to 3.*

**Remark 10** *Xiao and Ruan (see [10]) showed that either the number of infective individuals tends to zero as time evolves or the disease persists. Authors in [8, 20] proved that their epidemic model with saturated incidence rate, undergo a Bogdanov-Takens bifurcation of codimension 2. When we consider the incidence of a combination of the saturated incidence rate and a non-monotonic incidence, the codimension of Bogdanov-Takens bifurcation can be up to 3.*

## Conclusions

In this paper, by combing qualitative and bifurcation analyses, we study an SIS epidemic model with the incidence rate 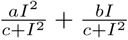, which can describe the inhibition effect from the behavioral change and interpret the “psychological” effect. In Section 2, we give a full-scale analysis for the types and stability of equilibria *E*_*i*_(*i* = 0, 1, 2, 3, 4, 5). We prove that system (2) can occur backward bifurcation and the backward bifurcation will disappear if *a* = 0. At equilibrium *E*_*i*_(*i* = 3, 5), degenerate Hopf bifurcation arises under certain conditions. When the critical condition ψ*_i_*(*i* = 3, 5) satisfied, we calculate the Liapunov value of the weak focus and obtain the value 2 for the maximal multiplicity of weak focus, indicating that there exist at most two limit cycles around *E*_*i*_(*i* = 3, 5). In Fig. 6, and Fig. 8, we give the phase portrait corresponding to equilibrium *E*_3_ and *E*_5_ about exhibiting a unique limit cycle and adding a new limit cycle after small perturbation of parameter λ and *c*. In Subsection 3.3, we proved that the model exhibits Bogdanov-Takens bifurcation of codimension 2 and codimension 3, under certain conditions. If parameter *a* = 0, the model can just have Bogdanov-Takens bifurcation of codimension 2, shown in [20].

In fact, we show that the model exhibits multi-stable states. This interesting phenomenon indicates that the initial states of an epidemic can determine the final states of an epidemic to extinct or not. Moreover, the periodical oscillation signify that the trend of the disease may be affected by the behavior of susceptible and the effect of psychology of the disease.

